# Opposing Roles of Cholesterol and Cholesterol Sulfate in Proton Transfer and Diffusion at Membrane Interfaces

**DOI:** 10.64898/2026.11.05.764480

**Authors:** Ambili Ramanthrikkovil Variyam, Mateusz Rzycki, Dominik Drabik, Nadav Amdursky

**Affiliations:** Schulich Faculty of Chemistry, Technion – Israel Institute of Technology, Haifa, 3200003, Israel; Department of Biomedical Engineering, Wroclaw University of Science and Technology, Wroclaw, 50-370, Poland; School of Mathematical and Physical Sciences, University of Sheffield, Sheffield S3 7HF, United Kingdom

**Keywords:** Cholesterol, Biological membranes, Proton diffusion, Proton transfer, Photoacids

## Abstract

Sterols are fundamental components of biological membranes that modulate membrane fluidity and organisation, yet their role in governing interfacial proton dynamics remains underexplored. Here, we elucidate how cholesterol (Ch) and cholesterol sulfate (ChS) regulate excited-state proton transfer (PT) and long-range proton diffusion (PD) using a complementary dual-approach experimental framework combined with molecular dynamics (MD) simulations. Using a membrane-anchored photoacid in small unilamellar vesicles, we demonstrate that Ch accelerates local PT dynamics in gel-phase phosphatidylcholine membranes by inducing a fluid, liquid-ordered phase. Surprisingly, ChS inhibits local PT despite its negative headgroup. MD simulations reveal this inhibition stems from deeper probe embedding within ChS-containing bilayers and the reduced basicity of sulfate relative to phosphate. Conversely, solid-state electrical measurements on supported lipid bilayers show a stark reversal for long-range proton conduction (lateral PD): ChS increases lateral proton conductance by 400-fold under controlled relative humidity, whereas Ch decreases conductivity. This divergence arises mainly because ChS supplies titratable sulfate charge carriers, whereas Ch dilutes key phosphate proton mediators that promote a percolating interfacial hydrogen-bonding network without providing charge carriers. Our findings demonstrate that sterols exert scale-dependent control over membrane PT/PD, revealing how headgroup chemistry, membrane fluidity, and hydration synergistically govern bioenergetic proton dynamics.

## Introduction

Proton diffusion (PD) along lipid membranes plays a fundamental role in cellular bioenergetics, particularly in processes such as ATP synthesis and cellular respiration (1, 2). It is well-established that membranes can support lateral PD, and an apparent energy barrier exists at the membrane-bulk interface for PT (3–10). Over the years, both experimental and computational studies have shown that factors such as membrane composition, lipid phase state, and lipid structure can significantly affect the proton barrier and PD along membranes (11–20).

The lipid composition of biological membranes is a complex assembly of a diverse range of lipids, including phospholipids, sphingomyelins, and sterols. Among sterols, cholesterol (Ch) is the most abundant in mammalian membranes and plays a crucial role in modulating membrane fluidity and permeability. In lipid membranes, Ch influences the membrane phase separation into liquid ordered (L_o_) and liquid disordered (L_d_) phases, with L_o_ having a higher degree of hydrocarbon chain order than L_d_ (21–25). Surprisingly, although Ch is an essential component of biological membranes, little is known about how it modulates PD dynamics at the membrane interface. Another biologically relevant sterol is cholesterol sulfate (ChS), which has a negatively charged sulfate group (**Figure 1**) and has been shown to contribute to membrane rigidity and fluidity (26). Unlike Ch, the biophysical properties of ChS are not well- explored. ChS is mainly found in the spermatozoan plasma membrane, stratum corneum, and erythrocyte membranes, and primarily serves stabilising functions in these cells (27–29). Given the important regulatory functions of sterols, we aim to investigate how Ch and ChS influence proton dynamics on lipid membranes. We hypothesise that by changing membrane order, surface hydration, electrostatic/dipole environment and lateral organisation, Ch/ChS can alter how protons diffuse along the membrane surface. Previous work found that Ch can reduce proton transport *across* the lipid bilayer (30). Nonetheless, our study provides the first direct correlation between lateral proton transfer (PT) and PD processes *on the membrane interface* as a function of Ch presence. We further compare the effect of Ch on the PT/PD processes with that of ChS, which, owing to its additional sulfate group, may modulate proton dynamics differently.

**Figure 1.**
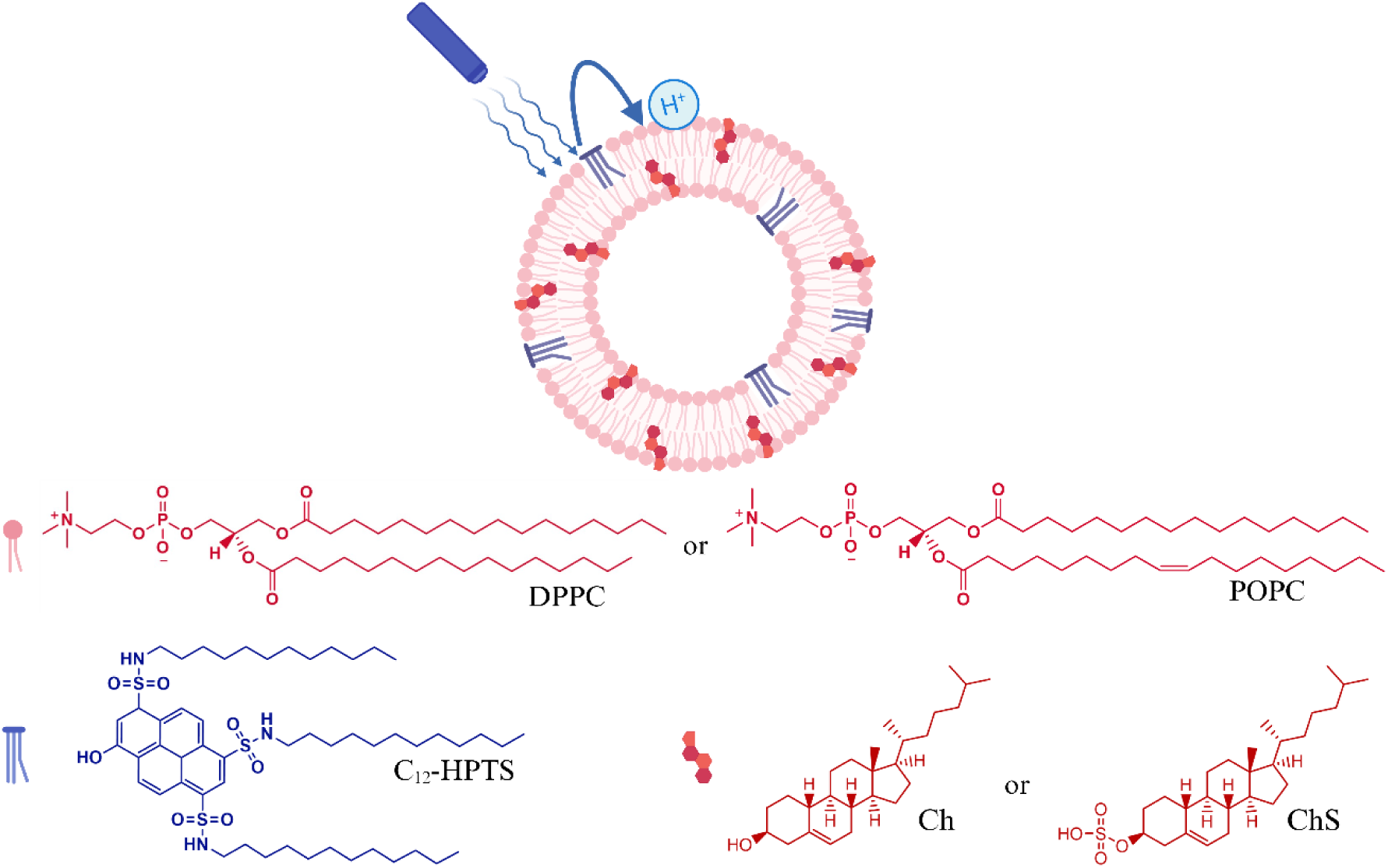
Schematic illustration of the ESPT process of C_12_-HPTS inside phospholipid membranes containing sterols. The chemical structures of DPPC, POPC, C_12_-HPTS, Ch and ChS are provided.

We use two distinct approaches to probe proton dynamics in sterol-containing lipid membranes. The first involves a fluorescent, membrane-anchored photoacid probe (13, 18–20). Photoacids are molecules that can undergo excited-state proton transfer (ESPT) upon illumination, owing to the drastic drop in pK_a_ values from the ground to the excited state. The dissociated proton from the photoacid can be accepted by the membrane, followed by several processes: PD on the membrane surface, escape to the bulk, or recombination with the now deprotonated photoacid. This approach allows us to monitor excited-state proton dynamics by measuring time-resolved fluorescence from the photoacid, since the protonated and deprotonated species emit at different wavelengths. In this approach, we use small unilamellar vesicles (SUVs) composed of lipid mixtures containing controlled amounts of Ch or ChS with phosphatidylcholine (PC). The second approach is based on electrical measurements on solid-state-supported lipid multi-bilayers (SLMBs) (31). This approach enables us to explore the long-range lateral proton conductivity on the membrane surface while controlling the exact amount of water molecules in a solid-state-like configuration by attenuating the relative humidity (RH) around the SLBs. Our work highlights the importance of fluidity, sterol composition, and water content in the ability of sterol-containing membranes to support PT and PD on their surface. We show that when the membrane acts as a proton acceptor in an SUV configuration, Ch enhances PT, whereas ChS surprisingly inhibits it. In contrast, in an SLMB configuration without bulk water, ChS, with its sulfate group, strongly enhances PD on the membrane surface, whereas Ch decreases it. We further explain our results using molecular dynamics (MD) simulations.

## Results and Discussions

To investigate the role of sterols in regulating proton dynamics of lipid membranes, we chose two different sterols: Ch and ChS. In the first part of our study, we followed PT processes on membranes containing these sterols. As model membranes, we used membranes composed primarily of 1,2-dipalmitoylphosphatidylcholine (DPPC) (in comparison to 1-palmitoyl-2- oleoyl-sn-glycero-3-phosphocholine (POPC)). The phospholipids were chosen due to their differences in main phase transition temperature (T_m_) at room temperature (RT). DPPC (T_m_ = 42 °C) is in the gel state at RT, whereas POPC (T_m_ = -2 °C) is in the fluid state. Although both Ch and ChS share a common hydrophobic backbone, Ch is a neutral molecule, whereas ChS possesses a negatively charged sulfate headgroup. To probe PT at the membrane interface, we used C_12_-HPTS (at a 1% mol ratio relative to lipids/sterols), a lipophilic derivative of pyranine that undergoes reversible proton dissociation upon light excitation (**Figure 1**).

The probe C_12_-HPTS exhibits two optical absorption bands corresponding to its protonated (ROH) and deprotonated (RO^-^) forms. Upon excitation of ROH, ESPT occurs:

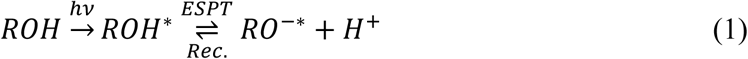

In the excited state, the proton dissociates (the ESPT process in eq. (1)), whereas the membrane acts as a proton acceptor. Subsequently, the released proton can either geminately recombine to reform ROH* (noted as Rec. in eq. (1)) or undergo PD from the photoacid, with membrane composition determining which process predominates. These post-ESPT processes are strongly influenced by the dimensionality (*d*) of PD along the membrane.

For an ESPT process, it is essential to ensure that we predominantly excite a protonated (ROH) photoacid. To confirm this, we performed pH-dependent UV-Vis absorption measurements to determine the ground-state pK_a_ of C_12_-HPTS in vesicles containing Ch and ChS (**Figure S1**), and we compared these values with the pK_a_ of the probe in pure POPC and DPPC membranes. The formation of homogeneous SUVs was verified by dynamic light scattering, which showed diameters of ∼100 nm across all lipid compositions. In the pK_a_ analysis, we worked with Ch or ChS mole ratio of 20% (compared to the concentration of lipids) within the membranes, a value that is in the range of common concentrations typically found in distinct biological membranes (32). Our results show that incorporating Ch into either DPPC or POPC vesicles leads to only a minor shift in the probe’s pK_a_. In contrast, adding ChS causes a more pronounced change, with the pK_a_ shifting to a higher value due to the negatively charged sulfate head group. The high sensitivity of the probe’s pK_a_ value to the neutral Ch vs the negatively charged ChS already suggests that the location of the probe’s -OH group is at the membrane-water interface. A similar trend was observed for the excited-state pK * of the probe across the different vesicles (**Table S1**), which was calculated using the Förster cycle (**Figure S2**).

## Steady-state fluorescence

Steady-state fluorescence provides an initial, straightforward validation of the stationary phase of the process and whether an ESPT process happened. The ratio between the protonated ROH* band (at 470 nm) and the deprotonated RO⁻* band (at 550 nm) can be used to extract the kinetic parameters of the ESPT process. A higher RO⁻*/ROH* emission ratio indicates faster deprotonation, and vice versa. As shown in **Figure 2a**, incorporating Ch (20% mol ratio) into DPPC significantly decreased the ROH* intensity (i.e., increased the RO⁻*/ROH* ratio), the first indication of enhanced ESPT of the probe within Ch-containing membranes. In our previous studies, we showed that negatively charged phosphatidic acid (PA) head groups accelerate ESPT from the probe (13, 18, 20). However, adding ChS (20% mol ratio) produced a surprising reverse trend in the normalised fluorescence spectrum, i.e., a lower RO⁻*/ROH* ratio, even though it has a negatively charged head group (**Figure 2a**).

**Figure 2.**
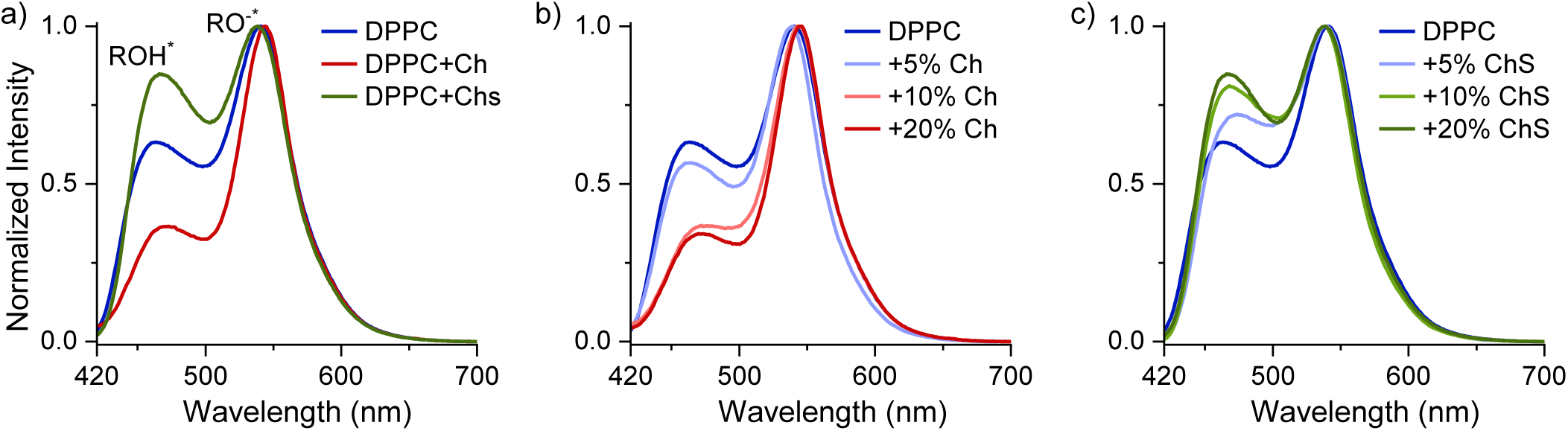
(a) Steady-state fluorescence of C_12_-HPTS inside DPPC, DPPC+Ch and DPPC+ChS. The concentrations of Ch and ChS are 20%. **(b)** and **(c)** Concentration dependence of Ch and ChS, respectively, within DPPC vesicles.

To understand this surprising trend, we have examined the membrane phase upon the addition of Ch/ChS. Using differential scanning calorimetry (DSC), the phase transition temperature (T_m_) of these membranes was determined (**Table S2**). Adding Ch clearly changes the phase of DPPC vesicles (T_m_ shifts from 43 to 40, 19, and 12°C with 5, 10, and 20% Ch, respectively) by interacting with the acyl chains of DPPC lipids. In contrast, adding ChS does not change the membrane T_m_ compared with that of pure DPPC (43°C). It is known that ChS increases the ordering of acyl chains of phospholipids, but much less than Ch (33). Accordingly, while DPPC and DPPC+ChS membranes are in their gel phase at RT, the DPPC+Ch membrane is in its fluid phase. As we showed in our previous study, a fluid-phase membrane serves as a better proton acceptor for the probe than a gel-phase membrane (20). The membrane phase can explain the observed change in fluorescence for the Ch-containing membrane, but it cannot explain why incorporating ChS slows the ESPT process (further discussion below).

To further pinpoint changes in the steady-state fluorescence at RT due to the presence of Ch or ChS, we performed a concentration-dependent study, varying the concentration of Ch or ChS in DPPC membranes from 5% to 20% (**Figure 2b** and **2c** for Ch and ChS, respectively). Adding 5% Ch to DPPC already induced a large change in steady-state fluorescence, increasing the RO⁻*/ROH* ratio. Adding 10% Ch induced another large change in steady-state fluorescence in the same direction, while increasing the concentration further to 20% did not induce a major change. Importantly, the T_m_ of both 10% and 20% Ch in DPPC is below RT (**Table S2**), and both are in their liquid phase. Thus, highlighting the role of membrane fluidity in the observed Ch-induced changes in PT dynamics at the membrane surface. With ChS addition, we also observe an immediate change in steady-state fluorescence upon adding 5% ChS to DPPC, but now with a decreased RO⁻*/ROH* ratio (**Figure 2c**). This decrease continues upon adding 10% and 20% ChS. As discussed, adding ChS does not change the membrane T_m_ (**Table S2**). Thus, this highlights the direct contribution of ChS in slowing the ESPT process.

### Time-resolved fluorescence

The ratio between protonated and deprotonated species can be used to extract the rate of PT and the rate of recombination as follows (34):

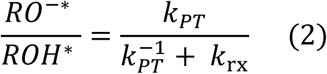

where *_kPT_* is the ESPT rate, *_k_*^−1^ is the proton recombination rate, and *k*_rx_ is the deactivation decay rate of *RO*^−∗^. The *k*_rx_ shares a similar value between pure DPPC membranes and vesicles containing Ch and ChS (0.2 ns^-1^) (**Figure S3**). To extract the kinetic parameters associated with PT on membranes, we used time-resolved fluorescence measurements.

In the time-resolved fluorescence measurements (**Figure 3a**), we monitored the decay of ROH* at 470 nm, which reflects three proton-dynamics processes associated with the membrane-bulk interface: ESPT occurs within the first few hundred picoseconds, followed by diffusion and recombination of the dissociated proton. In line with the steady-state measurements, we observed a faster fluorescence decay upon Ch addition. However, the differences manifested primarily after the first few hundred picoseconds associated with the ESPT process (the zoom- in area in **Figure 3a**), thus already highlighting that the observed changes are due to proton diffusion and recombination (discussion below). In contrast, ChS significantly slowed the initial ESPT process from the photoacid, as evidenced by a slower decay of ROH* (**Figure 3a**). To quantify the observed changes, we extracted *k_PT_* from the initial few hundred ps of the decay (using a three-exponential fit of the decay and considering the instrument response function) (**Table S3**). Compared to the ESPT rate of the C_12_-HPTS probe inside pure DPPC vesicles (3.5 ns^-1^), a significant decrease was observed upon incorporating ChS into the DPPC membrane, with a *k_PT_* dropping to 1.7 ns^-1^. In contrast, the incorporation of Ch resulted in a mild increase in the *k_PT_* to 3.9 ns^-1^.

**Figure 3.**
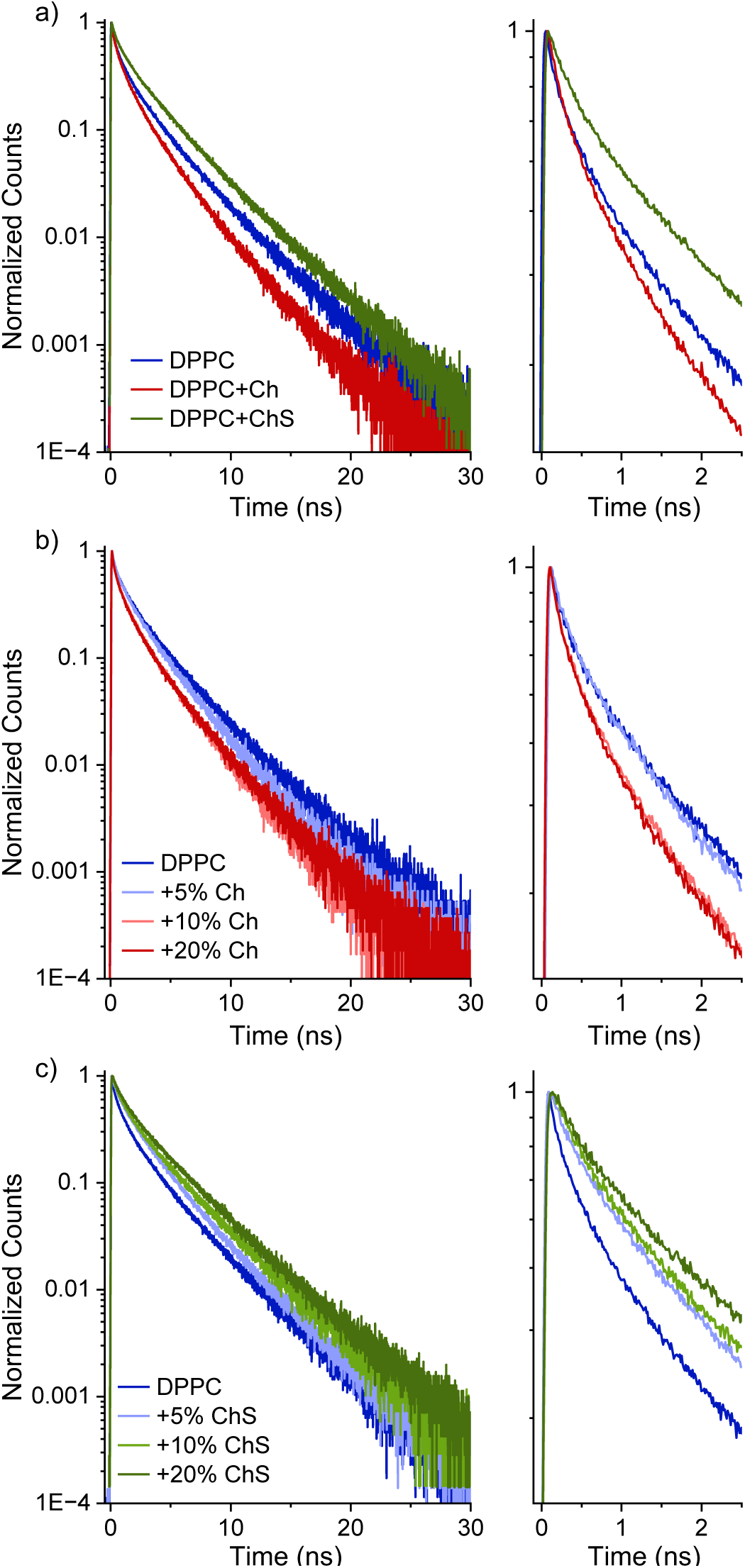
(a) Time-resolved fluorescence of C_12_-HPTS (at ROH* position) inside DPPC, DPPC+Ch and DPPC+ChS. The concentrations of Ch and ChS are 20%. **(b)** and **(c)** Concentration dependence of Ch and ChS, respectively, within DPPC vesicles. The right panels are a zoom-in of the first 2.5 ns of the decay.

The higher *k*_PT_ observed for Ch-containing DPPC membranes suggests that enhanced membrane fluidity can increase the ESPT from the photoacid. We observed a similar fluidity- dependent enhancement in *k*_PT_ in our previous study, where we increased temperature rather than composition to manipulate the membrane phase (20). Moreover, earlier reports have shown that cholesterol increases hydration at the headgroup region, which might well contribute to a higher *k*_PT_ (35–37). However, the rather similar value for DPPC highlights that the PC headgroup primarily dictates ESPT efficiency, whereas fluidity mainly influences the subsequent PD processes (see below). The next significant observation is that the sulfate headgroup impedes ESPT at the membrane interface compared with DPPC membranes, even though both are in their gel phase. The reduced ESPT rates in the presence of ChS are surprising. One explanation might be the reduced proton-accepting capability of sulfate compared with phosphate-based headgroups, since phosphates are stronger bases than sulfates. Therefore, membranes containing sulfate headgroups are poorer proton acceptors. Another explanation, revealed in our MD simulations (see below), is that ChS in DPPC shifts the C_12_- HPTS probe to a more buried location, which might also influence the probe’s initial ESPT.

As with the steady-state measurements, we have followed the time-resolved fluorescence at different concentrations of Ch (**Figure 3b**) or ChS (**Figure 3c**) in DPPC membranes from 5% to 20%. In Ch-containing membranes, the main changes manifest at longer (>1 ns) timescales after excitation, while the initial decay, which corresponds primarily to *k*_PT_, remains relatively similar across Ch concentrations (**Figure 3b** and **Table S3**). In contrast, for ChS-containing membranes, the measured fluorescence decay changes substantially even with the addition of 5% ChS (**Figure 3c**), primarily in the initial decay, highlighting the different *k*_PT_ across ChS concentrations (**Table S3**). As in the steady-state measurements, the overall trend between Ch and ChS is reversed: adding Ch at different concentrations results in faster decay, whereas adding ChS results in slower decay.

To distinguish between the roles of fluidity and headgroups, we compared our results with DPPC to POPC membranes. Owing to the lower saturation of the acyl chains in POPC, its membranes have a low T_m_ value (T_m_ = -2°C), hence, they are in their fluid phase regardless of the presence of Ch/ChS. Accordingly, we prepared POPC vesicles with the same concentration (20%) of Ch and ChS. As shown in both the steady-state measurements and the time-resolved measurements (**Figure S4**), adding Ch to the POPC SUVs yielded results nearly identical to those for POPC. These results suggest that Ch itself does not influence PT/PD dynamics under the same headgroup and fluidity conditions. However, as with our results for the DPPC membranes, incorporating ChS into POPC membranes also resulted in a lower RO⁻*/ROH* ratio and slower ROH* decay than in bare POPC membranes (**Figure S4**). These results highlight the effect of the sulfate group in inhibiting proton dynamics of the probe, regardless of lipid membrane fluidity.

In addition to the *k*_PT_ values that are extracted from the initial decay of ROH*, several other PD parameters can be obtained from the long-time fluorescence decay of C_12_-HPTS using a simplified diffusion model (34). Here, PD refers to proton movement around the excited-state probe. For example, on membranes, the PD dimensionality (*d*) equals 2, corresponding to a lateral PD, while *d* = 3 corresponds to both a lateral PD and proton escape from the surface to the bulk. As demonstrated in the model,(34) the long-lived tail (> 1 ns) of the photoacid’s ROH* decay follows a power-law behaviour, which is related to *d*:

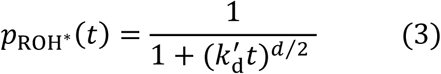

*p*_ROH_∗(*t*) is the ROH^∗^ population at time *t* and *k*^′^ is an effective rate of proton dissociation.

Accordingly, the next parameter that we will discuss is the *d* of PD (**Table S3**). The *d* value of pure DPPC vesicles is 1.9 at RT, where the membrane is in the gel state. As discussed in our previous study (20), a dimensionality below 2 implies PD pathways along membranes in the gel phase, whereas the transition to a liquid phase increases dimensionality. Consistent with our previous observations, we found that the *d* of Ch-incorporated DPPC vesicles is 2.7 (**Figure S5**), which is similar to the values obtained for liquid-phase PC membranes (such as for POPC) (18). This increase in *d* values indicates that membrane fluidity results in the ability of protons to escape from the membrane surface to the bulk. Hence, while the chemical nature of the phospholipid primarily influences the initial ESPT process (the similar initial decay between DPPC and DPPC+Ch in **Figure 3b**), the different membrane phase primarily influences the subsequent PD parameters. Incorporating ChS only mildly changed the *d* value compared with DPPC (a mild increase from 1.9 to 2.1, **Figure S5**). Since DPPC and DPPC+ChS share a similar T_m_, similar PD dimensionality associated with the gel phase is expected. From a structural perspective, the presence of ChS might interfere with the suggested PD pathways in the pure gel-phase DPPC membrane, thus resulting in attenuated *d*.

### Membrane phase-dependence of the ESPT process

All of the results above were conducted at RT, meaning that the DPPC and DPPC+ChS (at 20%) membranes were in their gel phase, while DPPC+Ch (at 20%) was in the liquid phase. In this section, we have varied the temperature and compared the steady-state (**Figure 4a**) and time-resolved (**Figure 4b**) fluorescence of C_12_-HPTS within the membranes, where all three membranes are either in their gel phase at a temperature (10°C) below their T_m_ or in their liquid phase at a temperature (50°C) above their T_m_. As shown in the spectra, the steady-state and time-resolved spectra of the probe in DPPC at the high temperature are highly similar to those in DPPC+Ch. This finding highlights that, while the membrane is in the liquid phase, incorporating Ch does not significantly attenuate PT processes on the membrane surface. Similar to the finding in RT, in the high-temperature regime, the incorporation of ChS also resulted in a smaller RO⁻*/ROH* ratio (higher ROH* peak) and a slower ROH* decay, with extracted *k*_PT_ values of 2.6 ns^-1^, compared to 4.7 and 4.9 ns^-1^ for DPPC and DPPC+Ch, respectively. In the low-temperature regime, the trend remains similar. The DPPC+ChS membrane exhibits the smallest RO⁻*/ROH* ratio and the slowest ROH* decay of all samples, with an extracted *k*_PT_ of 1.3 ns^-1^, compared to 3.1 and 3.2 ns^-1^ for DPPC and DPPC+Ch, respectively. These measurements further highlight that the membrane phase is the main factor dictating ESPT on the membrane surface in both DPPC and DPPC+Ch membranes, whereas incorporating ChS considerably attenuates ESPT in both phases. Below, we discuss the reasoning behind this.

**Figure 4.**
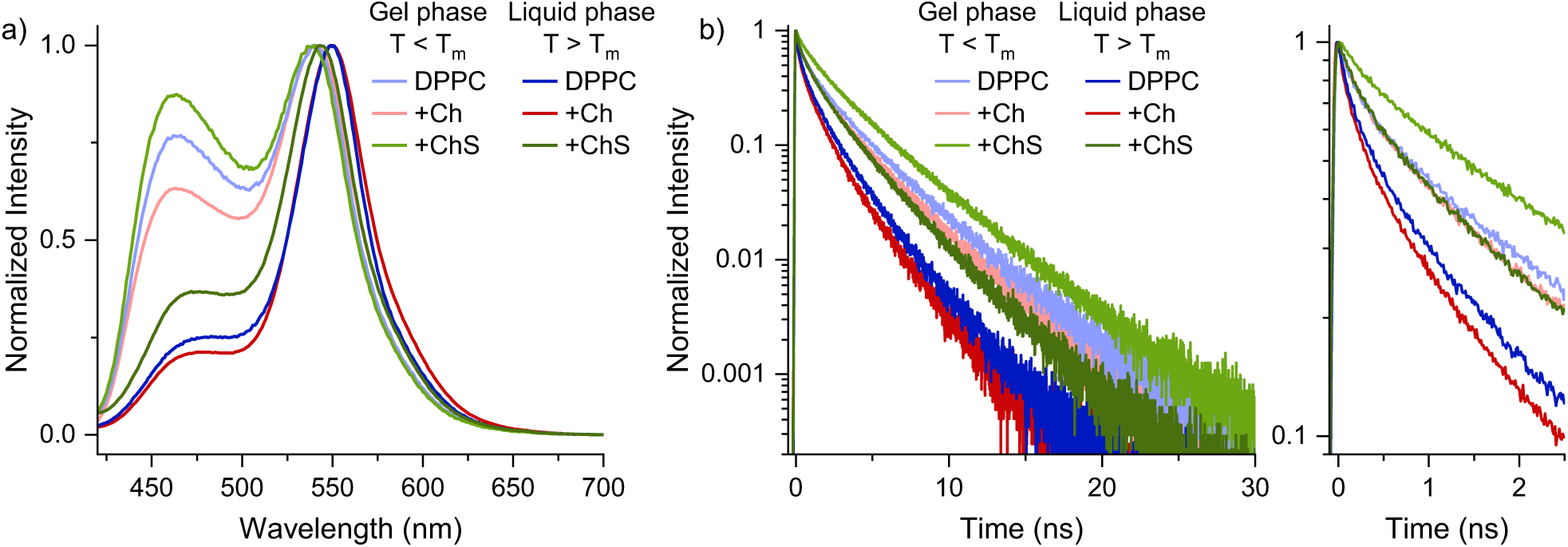
(a) Steady-state and (b) Time-resolved fluorescence of C12-HPTS in DPPC, DPPC+Ch, and DPPC+ChS taken at 10°C and 50°C, where the membranes are in their gel and liquid phases, respectively. The concentrations of Ch and ChS are 20%. The right panel in (b) is a zoom-in of the first 2.5 ns.

### Solid-state-like lateral proton conduction

To complement the ESPT part, which focuses on ultrafast PT from the probe to the membrane and short-range PD around the excited probe, we turned to measuring very long-range PD along the membranes using a solid-state supported lipid multi-bilayer (SLMB) approach. The solid- state environment allows for decoupling the contribution of bulk water molecules in PD by measuring the long-range proton conductivity along lipid membranes using electrochemical impedance spectroscopy (EIS). The SLMBs were prepared following the approach described in our previous study, resulting in a continuous layer of 4-5 multi-lipid bilayers, spanning between interdigitated electrodes (31). We fitted the EIS output to an equivalent circuit (**Figure S6**) to extract the proton resistance across the membrane-based 2-terminal device. In stark contrast to the previous part, we observed a striking 400-fold decrease in resistance (an increase in proton conductance) for ChS-containing DPPC membranes compared to DPPC SLMBs (**Table S4** and **Figure 5a**, as evidenced by a smaller semicircle in the EIS figure), indicating that ChS-containing DPPC membranes are more conductive. On the other hand, the Ch- containing membranes resulted in ∼2-fold increase in resistance compared to DPPC (**Table S4**). In general, solid-state proton conduction at an interface (the membrane interface in our work) is expected to depend on the surrounding RH, i.e., the number of water molecules adsorbed to its surface. More water molecules at the surface provide more opportunities for hydrogen-bond formation between water molecules and chemical moieties at the interface, resulting in higher protonic mobility (diffusion). The main advantage of the solid-state SLMBs configuration is our ability to attenuate the number of water molecules at the interface by changing the RH around the sample. In our RH-dependent measurements for all samples, we observed a sharp change in impedance (and thus proton resistance/conductance) when RH decreased from 80 to 50% (**Figure 5b**).

**Figure 5.**
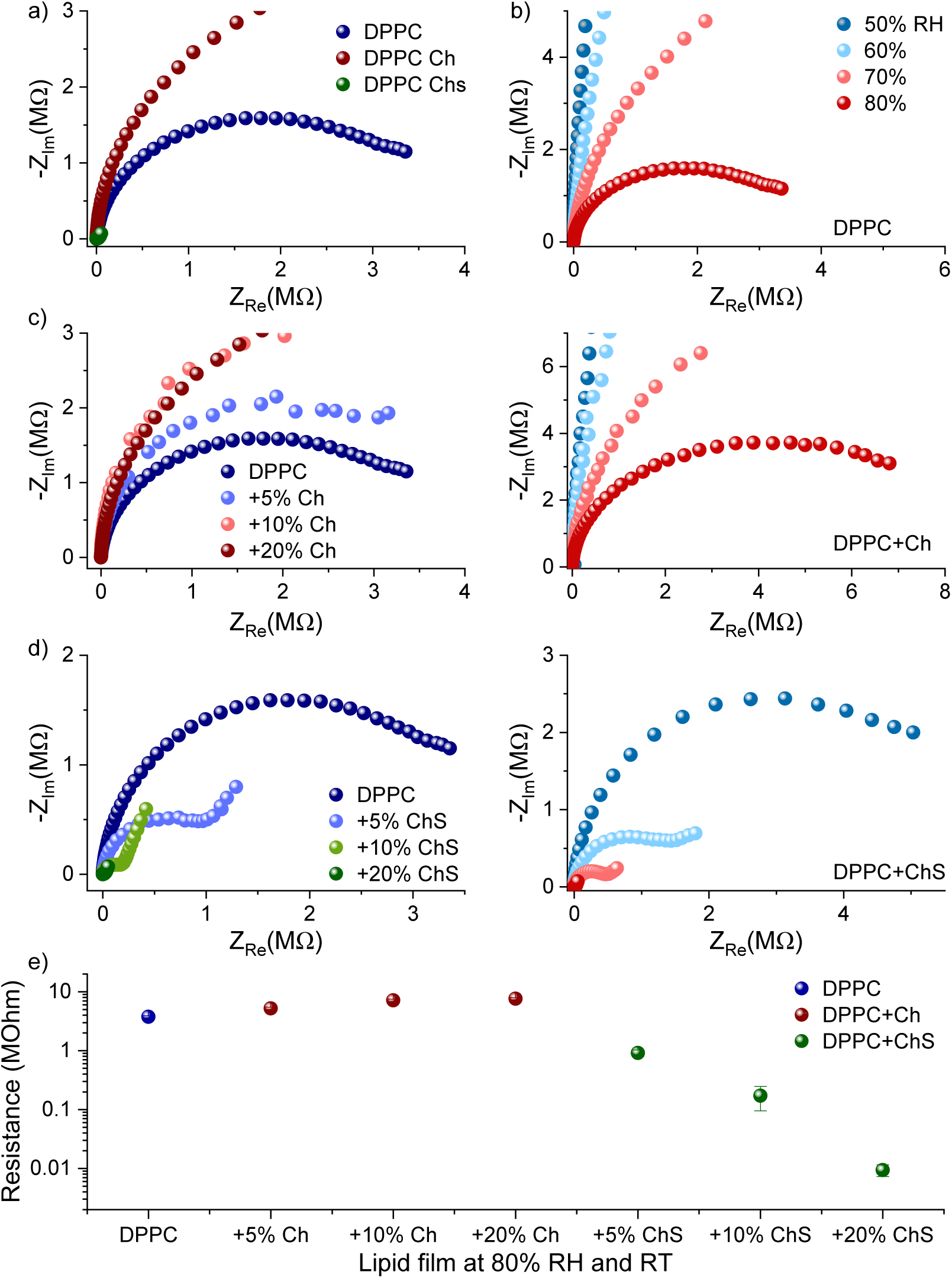
(a) EIS measurements (in the form of a Nyquist plot) of solid-state SLMBs composed of DPPC, DPPC+Ch, and DPPC+ChS at 80% RH and RT. The concentrations of Ch and ChS are 20%. **(b)** The RH-dependent EIS measurements of SLMBs in (a) (top – DPPC, middle – DPPC+Ch, bottom – DPPC+ChS). **(c)** and **(d)** Concentration dependence of Ch and ChS, respectively, within DPPC-based SLMBs. **(e)** A summary of the extracted resistance for the various SLMBs at 80% RH and RT.

In line with the ESPT measurements, we also followed the EIS across the membranes as a function of the concentration of either Ch or ChS (**Figure 5c** and **5d**, respectively). As shown by the change in extracted proton resistance (**Figure 5e** and **Table S4**), the amount of ChS within the membrane strongly affects the measured lateral proton conductance. On the other hand, increasing the concentration of Ch results in an increase in resistance, but the effect is considerably milder. These findings highlight the fundamental role of the ChS molecule in promoting lateral proton conduction at the membrane interface.

The contrast between the roles of Ch and ChS in ESPT from the probe to the membrane and in long-range lateral proton conduction highlights different factors associated with PT and PD on the membrane of biological membranes that are operating in different regimes. While membrane fluidity is the most important factor in the ESPT process and the membrane’s ability to accept protons, it is less important for long-range PD across membranes, as measured by proton conduction (31). The most important parameters for efficient long-range proton conduction are the number of protonic charge carriers and their mobility across the material. The additional sulfate group in ChS can increase the number of protonic charge carriers on the SLMB surface and facilitate better proton mobility at the membrane interface. Efficient proton mobility requires a percolating hydrogen-bond network at the membrane interface, comprising membrane functional groups (phosphates/sulfates/polar groups) and water molecules. This picture allows proton conduction via a Grotthuss-like mechanism. As a result, higher RH leads to more water molecules adsorbed at the membrane interface and better proton conductance. As shown below, ChS increases the propensity to form hydrogen bonds at the interface compared to Ch. In contrast, incorporating Ch does not contribute to either the hydrogen-bond network or additional protonic charge carriers. Hence, incorporating Ch depletes the concentration of DPPC phospholipids, which are important for proton conduction, resulting in lower measured proton conductance.

### Molecular dynamics simulations of the membranes

To better understand the contrasting results from Ch and ChS incorporation across our two measurement approaches, we performed MD simulations of the membranes used in this study.

Membrane properties and composition can dictate the efficiency of PT reactions on their surface (11, 13, 16, 18–20, 38). As shown in our experiments, ChS does not act as a common sterol and does not attenuate the membrane T_m_ upon incorporation. In our MD calculations, we observe a similar trend, where the addition of Ch (at the same 20% mol ratio) to a gel-phase DPPC induces an ordering of the membrane into a liquid-ordered phase, where the lipids retain a relatively high degree of acyl-chain ordering, while gaining more lateral mobility compared with the gel phase. This Ch-induced transition to a liquid-ordered phase increased membrane thickness (**Figure 6a**) and decreased the average area per lipid (**Figure 6b**) (39). Importantly, adding the C_12_-HPTS probe at the concentration used in this study does not substantially change these properties. As discussed, this Ch-induced transition to a liquid-ordered phase is the most likely reason for the higher ESPT rate from the probe to the membrane in DPPC+Ch membranes compared with other membranes. Since the incorporation of ChS has only a limited effect on the membrane properties (in terms of membrane thickness, **Figure 6a**, and area per lipid, **Figure 6b**), we would have expected similar PT properties from the probe to the membrane surface of DPPC and DPPC+ChS, but we found a surprisingly slower PT in the latter case. As we showed for several membrane systems (all in their fluid phase) (13), the headgroup of the C_12_-HPTS probe sits close to the membrane edge. From our MD simulations (**Figure 6c**), we find that this is the case only for the DPPC+Ch membrane, the only fluid membrane in our study. In the DPPC+Ch membrane, the probe (its OH group, i.e., the dissociated proton) sits merely 2.3±0.5 Å from the membrane edge. In contrast, in gel-phase DPPC membranes, the probe sits slightly deeper (4.3±0.6 Å from the membrane edge), which may further support slower PT from the probe in these membranes. Interestingly, in the DPPC+ChS (**Figure 6c**), the probe sits even deeper into the membrane (5.7±0.9 Å from the membrane edge). The probe location in the DPPC+ChS membrane may explain the slowest PT rate constant observed in this work.

**Figure 6.**
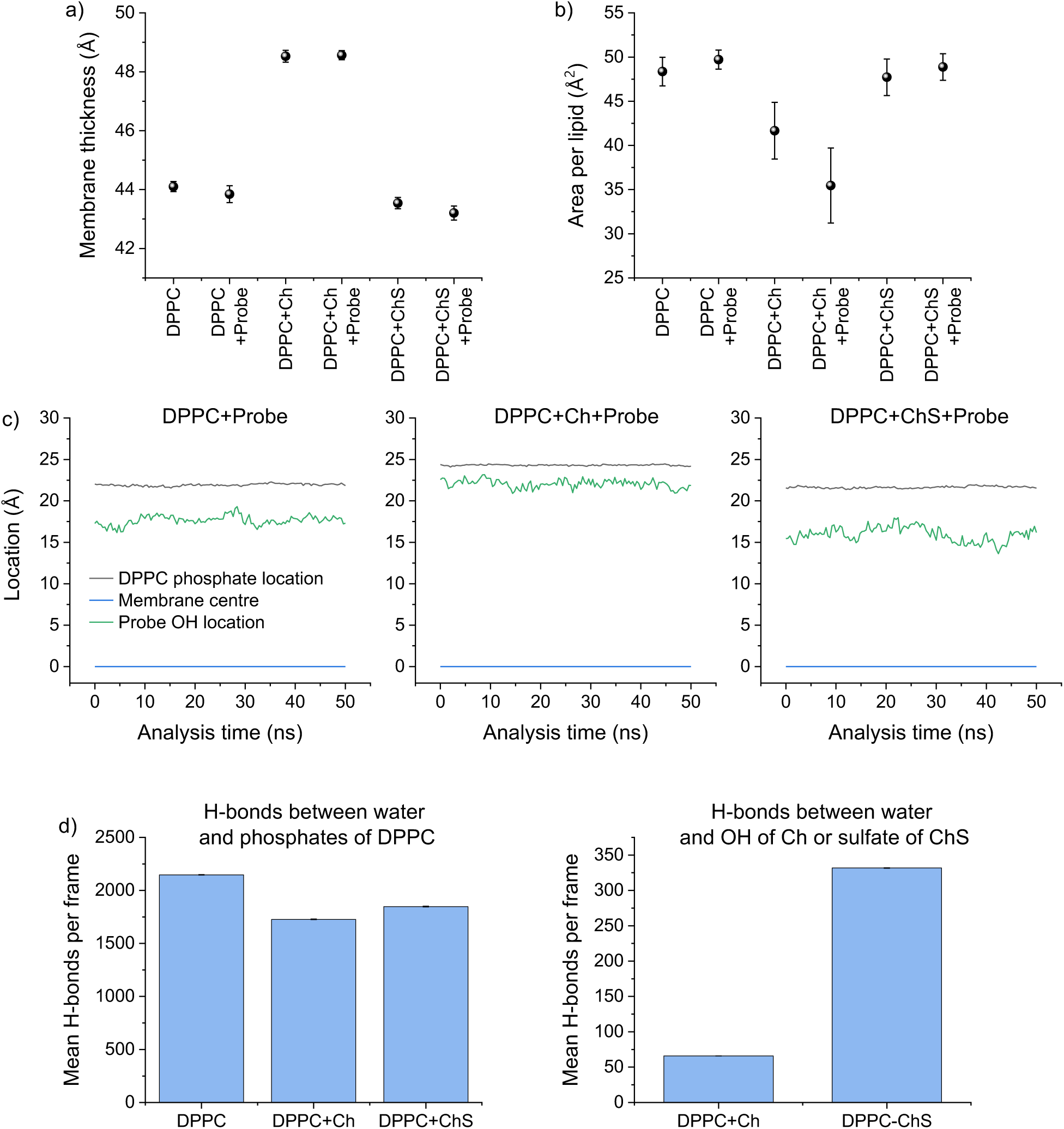
MD simulations of membrane properties. **(a)** Membrane thickness and **(b)** Area per lipid of DPPC, DPPC+Ch, and DPPC+ChS without or with the C_12_-HPTS probe. **(c)** The simulated location of the probe (the -OH group) within DPPC, DPPC+Ch, and DPPC+ChS membranes. **(d)** The calculated hydrogen bonds between water molecules and the phosphate groups of the phospholipids (left) and the -OH or sulfate group of Ch or ChS (respectively, right) at the different membrane interfaces.

As we moved from short-range PT steps from the probe to the membrane to the very long- range lateral PD processes observed in our electrical measurements, we discussed two key parameters: protonic charge carriers and proton mobility. The concentration of protonic charge carriers is mainly dictated by the presence of titratable (ionizable) groups. In our work, the most accessible titratable group is the sulfate group of ChS. For proton mobility, one of the most crucial parameters is the hydrogen bonding at the membrane interface. The more hydrogen bonds that can form at this interface, the better the proton mobility. In our MD calculations, we have calculated the number of hydrogen bonds (per frame of the MD simulation) between nearby water molecules and several groups in the membrane: the phosphate group of the DPPC phospholipids, the sulfate group of ChS, and the OH group of Ch (**Figure 6d**). By far, the number of hydrogen bonds to the phosphate group is larger than the number of hydrogen bonds to either ChS or Ch. The number of hydrogen bonds to the sulfate group of ChS is substantially larger than those to the OH group of Ch, suggesting higher mobility on the surface of DPPC+ChS than on DPPC+Ch. Nonetheless, in absolute terms, this number of hydrogen bonds is significantly lower than to phosphate groups. Importantly, both Ch and ChS deplete the concentration of DPPC in the system, resulting in fewer hydrogen bonds to phosphate in membranes containing ChS or Ch, whereas the depletion is larger for DPPC+Ch than for DPPC+ChS (**Figure 6d**). Accordingly, we may conclude that the large increase in measured proton conduction in the DPPC+ChS membrane is primarily due to the higher protonic charge-carrier concentration provided by the sulfate group of ChS at the membrane interface. Compared with DPPC+Ch, we expect proton mobility to be greater in DPPC+ChS, further widening the difference between them. Although we previously established that a liquid-phase membrane is a better proton conductor than a gel-phase membrane (31), the lowest number of hydrogen bonds at the membrane interface of DPPC+Ch is probably the reason for the highest resistance measured for this membrane.

## Concluding Remarks

In summary, we provide a comprehensive investigation into how sterols modulate proton dynamics at biological membrane-water interfaces. By integrating time-resolved fluorescence spectroscopy of a membrane-bound photoacid, solid-state EIS measurements of proton conduction, and atomistic MD simulations, we demonstrate that Ch and ChS have fundamentally distinct and opposing effects on short-range PT versus long-range PD. At short range, the measured ESPT between the tethered photoacid and the membrane surface shows that the membrane’s physical state and the probe microenvironment dominate ESPT. The incorporation of Ch into gel-phase DPPC bilayers accelerates the ESPT rate constant (*k_PT_*), which is primarily driven by Ch-induced transition from the gel phase to a liquid-ordered phase.

Conversely, in inherently fluid membranes (such as POPC), Ch incorporation induces minimal change in local PT dynamics. In contrast, the negatively charged sterol ChS produces an unexpected slowdown in local ESPT across both DPPC and POPC membranes. Ch incorporation and the transition to a liquid phase also allow surface-to-bulk proton escape, as indicated by an increase in PD dimensionality around the probe to values closer to 3, calculated from the longer fluorescence-decay timescales. The poor ESPT from the probe in the ChS-containing membranes is explained by two structural origins: 1) Sulfate headgroups are weaker proton acceptors than phosphate groups, making ChS-rich domains intrinsic thermodynamic sinks that impede local proton uptake. 2) Our MD simulations indicate that the photoacid headgroup sinks deeper into ChS-containing bilayers compared to Ch-containing membranes and even pure DPPC membranes, creating a spatial barrier that hinders initial proton release to the interface.

In stark contrast to our ESPT study, long-range lateral proton conduction across supported lipid bilayers reveals a complete reversal of sterol efficacy. EIS measurements under controlled RH show that ChS nearly doubles long-range lateral proton conductivity compared to pure DPPC, whereas Ch modestly decreases it. Overall, long-range PD (lateral proton conduction) depends on protonic charge-carrier density and a continuous, percolating hydrogen-bond network across the membrane surface. The sulfate group of ChS provides additional titratable, ionisable charge carriers and effectively retains interfacial water molecules, facilitating efficient Grotthuss-like proton hopping even under reduced hydration. Conversely, neutral Ch lacks ionisable headgroups and displaces native phospholipid headgroups, thereby diluting the interfacial phosphate network essential for lateral proton migration without supplying alternative charge carriers.

Ultimately, these findings emphasise that sterol modulation of interfacial proton dynamics cannot be generalised by membrane fluidity or headgroup charge alone. Instead, proton behaviour is inherently scale-dependent: short-range local PT depends on headgroup properties, local hydration, probe positioning, and phase state, whereas long-range lateral PD is governed by charge carrier density and structural continuity of the interfacial hydrogen-bonding network. Given the distinct subcellular localisation of Ch and ChS in specialised biological membranes, such as spermatozoa, erythrocytes, and the stratum corneum, these contrasting physical mechanisms offer new bioenergetic and physiological insights into how biological systems fine-tune lateral proton microcircuits and membrane bioenergetics through localised lipid composition.

## Experimental Section

### Vesicle preparation

All the lipids were purchased from Avanti Polar Lipids. A 2 mM lipid solution was prepared in chloroform with a probe-to-lipid ratio of 1:100. The solvent was then evaporated to form a dry lipid film. This film was rehydrated using a 5 mM phosphate buffer at pH 7.4, and the mixture was extruded through a polycarbonate membrane (T&T Scientific) to obtain a homogeneous solution.

### Steady-state spectroscopic measurements

Steady-state UV-visible absorption and fluorescence emission experiments were measured using an Agilent Cary 60 spectrometer and an FS5 spectrometer (Edinburg Instruments), respectively. Dynamic light scattering (DLS) experiments were carried out using a Zetasizer Nano ZS (Malvern Instruments). Differential scanning calorimetry (DSC) measurements were performed using a DSC-1 (Mettler Toledo).

### Time-correlated single photon counting spectroscopy

The fluorescence lifetime of the probe inside different vesicles was measured using a CHIMERA spectrometer (Light Conversion). The samples were excited with a 400 nm pulse, and the time-resolved signal was collected using a hybrid detector (Becker & Hickl, HPM-100-07). The laser system used for these measurements is a PHAROS Yb-based laser (Light Conversion), which delivers 190 fs pulses at 1030 nm, operates at 1 MHz, and provides up to 10 mJ pulse energy. This beam was sent into an ORPHEUS optical amplifier to generate the required wavelengths through harmonic generation.

### Electrochemical impedance spectroscopy

Proton conductivity measurements were carried out using an impedance/gain-phase analyzer (MTZ-35, Bio-Logic). Lipid samples were spin- coated on the prepared electrodes: interdigitated gold electrodes with a 70 µm distance between electrodes. The samples were prepared according to the reported procedure.(31) The electrodes were made to be in contact using a probe station micromanipulator. A 50 mV AC bias was applied during the measurements, and a frequency range of 10 MHz to 10 Hz was used for the experiments. The conductivity of the films was calculated using the following equation: G = σA/l, where G is the conductance (as extracted using the equivalent circuit), σ is the conductivity, A is the cross-sectional area of the lipid film (A = thickness of film × width of the film), and l is the distance between two electrodes.

### Molecular modelling

#### System Construction and Parametrisation

Membrane models were constructed using CHARMM-GUI and further supported by custom Python scripts.(40, 41) Three lipid compositions were investigated: pure DPPC, DPPC:Ch at a 4:1 molar ratio, and DPPC:ChS at a 4:1 molar ratio. Each membrane contained 500 lipid molecules and was hydrated with 50 water molecules per lipid. NaCl was added to obtain a final concentration of 5 mM. In addition to the probe-free systems, replicate membrane models containing 1.5% of the C12-HPTS probe were prepared to investigate its behaviour in the different lipid environments. The parameters for C_12_-HPTS and ChS were initially generated using the CHARMM General Force Field (CGenFF).(42) For C_12_-HPTS, the resulting CGenFF description was used for subsequent simulations. Although the initial CGenFF assignment for ChS was associated with low penalty scores, its partial atomic charges were further refined using the standard Force Field Toolkit (ffTK)(43) charge-optimisation procedure based on quantum-mechanical water-solute interaction sites at the Hartree-Fock level of theory (HF/6-31G(d)). Particular attention was given to refining the electrostatic description of the linker region connecting the cholesterol-derived scaffold and the sulfate moiety to ensure consistency with the corresponding chemical fragments already parametrised within the CHARMM force field.(44)

#### Molecular Dynamics Simulations

All molecular dynamics simulations were performed using GROMACS.(45) Before production simulations, all membrane systems were subjected to energy minimisation and equilibration following the protocol employed in our previous study.(46, 47) Briefly, equilibration comprised 250 ps in the NVT ensemble using a 1 fs integration time step, followed by NPT equilibration for 250 ps with a 1 fs time step and then 5 ns with a 2 fs time step. Positional and dihedral restraints were gradually reduced throughout the equilibration procedure. MD simulations were performed for 300 ns using a 2 fs integration time step. The temperature was maintained at 295.15 K using the velocity-rescaling thermostat with a coupling constant of 1 ps. Pressure was maintained at 1 bar using the C-rescale barostat with semi-isotropic coupling, a coupling constant of 5 ps, and a compressibility of (4.5×10^-5^ bar^-1^). Bonds involving hydrogen atoms were constrained using the LINCS algorithm.

#### Trajectory Analysis

The final 50 ns of each production trajectory were used for quantitative analysis. Trajectory processing and analysis were performed using custom Python scripts together with the MDAnalysis library.(48) The procedures used to determine membrane thickness (MT), area per lipid (APL) and density profiles followed our previously established methodology.(46) Hydrogen bonds were analysed using a custom Python script based on the MDAnalysis library. A hydrogen bond was assigned when the donor-hydrogen distance was ≤1.2 Å, the donor-acceptor distance was ≤3.5 Å, and the donor-hydrogen-acceptor angle was ≥150°.

## Acknowledgements

N.A thanks the VolkswagenStiftung (number ZN5614), BBSRC standard grant (UKRI2988), and UKRI awards, Horizon Europe guarantee scheme grant (number EP/Z000459/1) for financial support.

## References

1. A. Y. Mulkidjanian, J. Heberle, D. A. Cherepanov, Protons @ interfaces: Implications for biological energy conversion. Biochim. Biophys. Acta-Bioenerg. 1757, 913–930 (2006).

2. R. J. P. Williams, Proton Circuits in Biological Energy Interconversions. Annu. Rev. Biophys. Biophys. Chem. 17, 71–97 (1988).

3. J. Heberle, J. Riesle, G. Thiedemann, D. Oesterhelt, N. A. Dencher, Proton migration along the membrane surface and retarded surface to bulk transfer. Nature 370, 379–382 (1994).

4. J. Heberle, N. A. Dencher, Surface-bound optical probes monitor protein translocation and surface potential changes during the bacteriorhodopsin photocycle. Proc. Natl. Acad. Sci. 89, 5996–6000 (1992).

5. U. Alexiev, R. Mollaaghababa, P. Scherrer, H. G. Khorana, M. P. Heyn, Rapid long- range proton diffusion along the surface of the purple membrane and delayed proton transfer into the bulk. Proc. Natl. Acad. Sci. 92, 372–376 (1995).

6. O. A. Gopta, D. A. Cherepanov, W. Junge, A. Y. Mulkidjanian, Proton transfer from the bulk to the bound ubiquinone Q(B) of the reaction center in chromatophores of Rhodobacter sphaeroides: Retarded conveyance by neutral water. Proc. Natl. Acad. Sci. 96, 13159–13164 (1999).

7. D. A. Cherepanov, W. Junge, A. Y. Mulkidjanian, Proton transfer dynamics at the membrane/water interface: dependence on the fixed and mobile pH buffers, on the size and form of membrane particles, and on the interfacial potential barrier. Biophys. J. 86, 665–680 (2004).

8. C. Zhang et al., Water at hydrophobic interfaces delays proton surface-to-bulk transfer and provides a pathway for lateral proton diffusion. Proc. Natl. Acad. Sci. 109, 9744– 9749 (2012).

9. Maarten G. Wolf, H. Grubmüller, G. Groenhof, Anomalous Surface Diffusion of Protons on Lipid Membranes. Biophys. J. 107, 76–87 (2014).

10. E. Weichselbaum et al., Origin of proton affinity to membrane/water interfaces. Sci. Rep. 7, 4553 (2017).

11. A. Springer, V. Hagen, D. A. Cherepanov, Y. N. Antonenko, P. Pohl, Protons migrate along interfacial water without significant contributions from jumps between ionizable groups on the membrane surface. Proc. Natl. Acad. Sci. 108, 14461–14466 (2011).

12. M. Brändén, T. Sandén, P. Brzezinski, J. Widengren, Localized proton microcircuits at the biological membrane–water interface. Proc. Natl. Acad. Sci. 103, 19766–19770 (2006).

13. N. Amdursky, Y. Lin, N. Aho, G. Groenhof, Exploring fast proton transfer events associated with lateral proton diffusion on the surface of membranes. Proc. Natl. Acad. Sci. 116, 2443–2451 (2019).

14. E. Weichselbaum, T. Galimzyanov, O. V. Batishchev, S. A. Akimov, P. Pohl, Proton Migration on Top of Charged Membranes. Biomolecules 13, 352 (2023).

15. E. S. Medvedev, A. A. Stuchebrukhov, Kinetics of proton diffusion in the regimes of fast and slow exchange between the membrane surface and the bulk solution. J. Math. Biol. 52, 209–234 (2006).

16. T. Sandén, L. Salomonsson, P. Brzezinski, J. Widengren, Surface-coupled proton exchange of a membrane-bound proton acceptor. Proc. Natl. Acad. Sci. 107, 4129–4134 (2010).

17. N. Agmon, M. Gutman, Proton fronts on membranes. Nat. Chem. 3, 840–842 (2011).

18. A. R. Variyam et al., Proton diffusion on the surface of mixed lipid membranes highlights the role of membrane composition. Biophys. J. 123, 4200–4210 (2024).

19. A. R. Variyam, M. Mencía, N. Amdursky, The role of ester-versus ether-linked phospholipids in the ability of biological membranes to accept protons and support proton diffusion. Biophys. J. 124, 2677–2685 (2025).

20. A. R. Variyam et al., The role of lipid phase and temperature in proton barrier and proton migration on biological membranes. Chem. Sci. 16, 5194–5204 (2025).

21. T. T. Bui, K. Suga, H. Umakoshi, Ergosterol-induced ordered phase in ternary lipid mixture systems of unsaturated and saturated phospholipid membranes. J. Phys. Chem. B 123, 6161–6168 (2019).

22. T. Róg, I. Vattulainen, Cholesterol, sphingolipids, and glycolipids: what do we know about their role in raft-like membranes? Chem. Phys. Lipids 184, 82–104 (2014).

23. S. L. Veatch, S. L. Keller, Organization in lipid membranes containing cholesterol. Phys. Rev. Lett. 89, 268101 (2002).

24. M. Cebecauer et al., Membrane lipid nanodomains. Chem. Rev. 118, 11259–11297 (2018).

25. F. Schmid, Physical mechanisms of micro-and nanodomain formation in multicomponent lipid membranes. *Biochim. Biophys. Acta*, Biomembr. 1859, 509–528 (2017).

26. A. Reis, M. J. Sarmento, M. Ferreira, P. Gameiro, V. de Freitas, Cholesterol Sulfate in Biological Membranes: A Biophysical Study in Cholesterol-Poor and Cholesterol-Rich Biomimetic Models. Membranes 15, 159 (2025).

27. G. Bleau, G. Lalumière, A. Chapdelaine, K. D. Roberts, Red cell surface structure: stabilization by cholesterol sulfate as evidenced by scanning electron microscopy. *Biochim. Biophys. Acta*, Biomembr. 375, 220–223 (1975).

28. P. Elias et al., Stratum corneum lipids in disorders of cornification. Steroid sulfatase and cholesterol sulfate in normal desquamation and the pathogenesis of recessive X- linked ichthyosis. J. Clin. Investig. 74, 1414–1421 (1984).

29. G. Lalumiere, G. Bleau, A. Chapdelaine, K. Roberts, Cholesteryl sulfate and sterol sulfatase in the human reproductive tract. Steroids 27, 247–260 (1976).

30. Ira, G. Krishnamoorthy, Probing the link between proton transport and water content in lipid membranes. J. Phys. Chem. B 105, 1484–1488 (2001).

31. A. Ramanthrikkovil Variyam, M. Stolov, J. Feng, N. Amdursky, Solid-State Molecular Protonics Devices of Solid-Supported Biological Membranes Reveal the Mechanism of Long-Range Lateral Proton Transport. ACS Nano 18, 5101–5112 (2024).

32. V. Schoop, A. Martello, E. R. Eden, D. Höglinger, Cellular cholesterol and how to find it. *Biochim. Biophys. Acta*, Mol. Cell. Biol. Lipids 1866, 158989 (2021).

33. A. M. Smondyrev, M. L. Berkowitz, Molecular dynamics simulation of dipalmitoylphosphatidylcholine membrane with cholesterol sulfate. Biophys. J. 78, 1672–1680 (2000).

34. A. A. Stuchebrukhov, A. R. Variyam, N. Amdursky, Using Proton Geminate Recombination as a Probe of Proton Migration on Biological Membranes. J. Phys. Chem. B 126, 6026–6038 (2022).

35. N. Kučerka, J. Pencer, M.-P. Nieh, J. Katsaras, Influence of cholesterol on the bilayer properties of monounsaturated phosphatidylcholine unilamellar vesicles. Eur. Phys. J. E. 23, 247–254 (2007).

36. W. K. Subczynski, A. Wisniewska, J.-J. Yin, J. S. Hyde, A. Kusumi, Hydrophobic barriers of lipid bilayer membranes formed by reduction of water penetration by alkyl chain unsaturation and cholesterol. Biochemistry 33, 7670–7681 (1994).

37. D. Marsh, Polarity and permeation profiles in lipid membranes. Proc. Natl. Acad. Sci. 98, 7777–7782 (2001).

38. J. Zhang, P. R. Unwin, Proton Diffusion at Phospholipid Assemblies. J. Am. Chem. Soc. 124, 2379–2383 (2002).

39. D. Drabik, G. Chodaczek, S. Kraszewski, M. Langner, Mechanical properties determination of DMPC, DPPC, DSPC, and HSPC solid-ordered bilayers. Langmuir 36, 3826 (2020).

40. S. Jo, J. B. Lim, J. B. Klauda, W. Im, CHARMM-GUI Membrane Builder for mixed bilayers and its application to yeast membranes. Biophys. J. 97, 50–58 (2009).

41. J. Lee et al., CHARMM-GUI input generator for NAMD, GROMACS, AMBER, OpenMM, and CHARMM/OpenMM simulations using the CHARMM36 additive force field. J. Chem. Theory Comput. 12, 405–413 (2016).

42. K. Vanommeslaeghe et al., CHARMM general force field: A force field for drug-like molecules compatible with the CHARMM all-atom additive biological force fields. J. Comput. Chem. 31, 671–690 (2010).

43. C. G. Mayne, J. Saam, K. Schulten, E. Tajkhorshid, J. C. Gumbart, Rapid parameterization of small molecules using the force field toolkit. J. Comput. Chem. 34, 2757–2770 (2013).

44. J. B. Klauda et al., Update of the CHARMM all-atom additive force field for lipids: validation on six lipid types. J. Phys. Chem. B 114, 7830 (2010).

45. M. J. Abraham et al., GROMACS: High performance molecular simulations through multi-level parallelism from laptops to supercomputers. SoftwareX 1, 19–25 (2015).

46. M. Rzycki, D. Drabik, Multifaceted activity of fabimycin: Insights from molecular dynamics studies on bacterial membrane models. J. Chem. Inf. Model. 64, 4204 (2024).

47. M. Rzycki, D. Drabik, K. Szostak-Paluch, B. Hanus-Lorenz, S. Kraszewski, Unraveling the mechanism of octenidine and chlorhexidine on membranes: Does electrostatics matter? Biophys. J. 120, 3392–3408 (2021).

48. R. Gowers et al., MDAnalysis: A Python Package for the Rapid Analysis of Molecular Dynamics Simulations. SciPy, 98–105 (2016).

