## Supporting Tables and Figures for "Opposing Roles of Cholesterol and Cholesterol Sulfate in Proton Transfer and Diffusion at Membrane Interfaces"

| Lipid membrane | $\lambda_{ROH}$ (abs), nm | $\lambda_{RO^-}$ (abs), nm | $\lambda_{ROH}$ (em), nm | $\lambda_{RO^-}$ (em), nm | $pK_a$ | $\Delta pK_a$ | $pK_a$* |
|---|---|---|---|---|---|---|---|
| DPPC | 420 | 499 | 461 | 540 | 9 | 7.3 | 1.7 |
| DPPC+Ch | 422 | 503 | 473 | 540 | 8.9 | 7.7 | 1.2 |
| DPPC+ChS | 421 | 499 | 466 | 539 | 9.6 | 7.6 | 2.0 |
| POPC | 422 | 501 | 476 | 555 | 8.1 | 7.2 | 0.9 |
| POPC+Ch | 423 | 503 | 470 | 554 | 8.5 | 7.0 | 1.5 |
| POPC+ChS | 422 | 501 | 473 | 544 | 9.2 | 7.7 | 1.5 |

**Table S1.** The ground and excited state $pK_a$ of photoacid inside the liposomes along with absorption and emission maxima for both ROH and $RO^-$ forms. The sterol concentration of vesicles id 20 mol%.

| Lipid membrane | $T_m$ (°C) |
|---|---|
| DPPC | 43 |
| DPPC + 5% Ch | 40 |
| DPPC + 10% Ch | 19 |
| DPPC + 20% Ch | 12 |
| DPPC + 5-20 ChS | 43 |

**Table S2.** Melting points of different concentrations of Ch containing DPPC membranes calculated from DSC.

| **Lipid membrane** | **$k_{PT}$ (ns$^{-1}$)** | ***d*** |
|---|---|---|
| DPPC | 3.5 | 1.9 |
| DPPC + 5% Ch | 3.5 | 2.4 |
| DPPC + 10% Ch | 3.8 | 2.7 |
| DPPC + 20% Ch | 3.9 | 2.7 |
| DPPC + 5% ChS | 2.8 | 2.4 |
| DPPC + 10% ChS | 2.2 | 2.5 |
| DPPC + 20% ChS | 1.7 | 2.1 |

**Table S3.** The extracted $k_{PT}$ values (from the initial decay $ROH^*$) and dimensionality (*d*) from eq.(3) in the main text for the time-resolved measurements at RT.

| **SLMB** | **Resistance (Ω)** |
|---|---|
| DPPC | $3.8\times10^6$ |
| DPPC + 5% Ch | $5.3\times10^6$ |
| DPPC + 10% Ch | $7.1\times10^6$ |
| DPPC + 20% Ch | $7.6\times10^6$ |
| DPPC + 5% ChS | $9.1\times10^5$ |
| DPPC + 10% ChS | $1.7\times10^5$ |
| DPPC + 20% ChS | $9.4\times10^3$ |

**Table S4.** The resistance of SLMBs at different lipid compositions measured using EIS at RT and 80% RH.

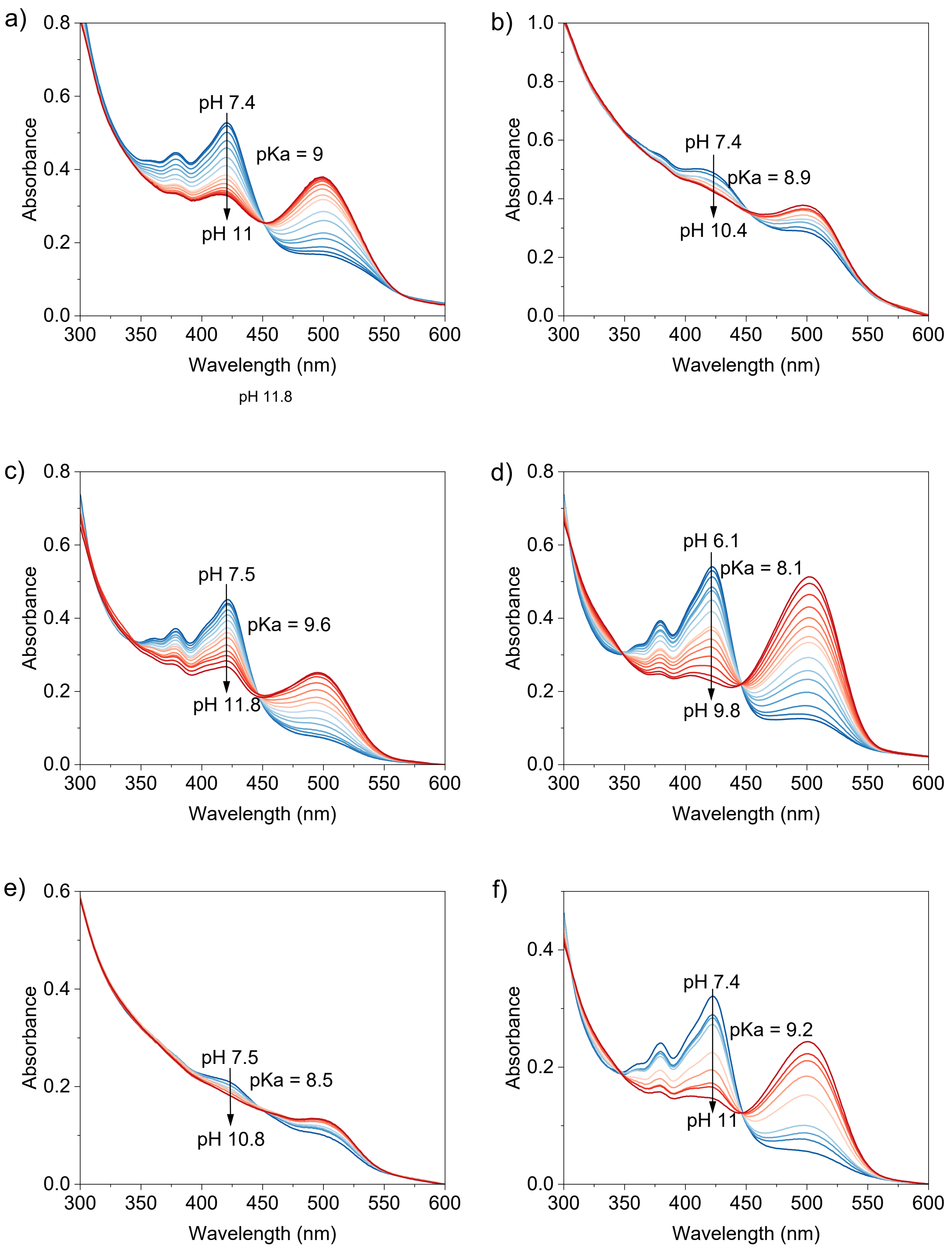


**Figure S1.** pH titration of $C_{12}$-HPTS inside (a) DPPC, (b) DPPC+Ch, (c) DPPC+ChS, (d) POPC, (e) POPC+Ch, (f) POPC+ChS SUVs using UV- Vis absorption spectroscopy. The sterol concentration is 20 mol%.

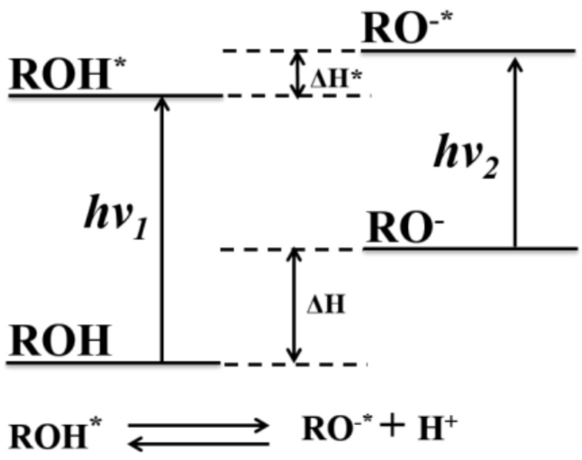


**Figure S2.** Förster cycle of photoacids.

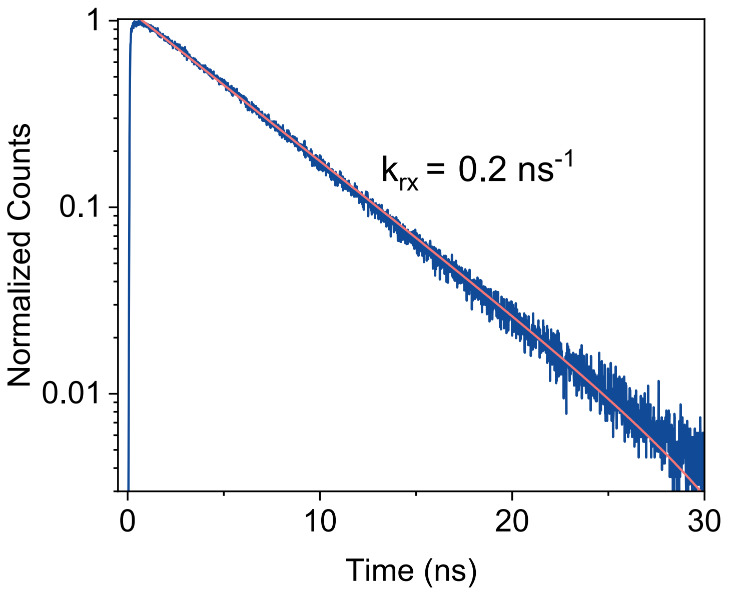


**Figure S3.** Rate of fluorescence radiative decay ($k_{rx}$) of photoacid determined from the $RO^{-*}$ decay. This value is the same for all membrane compositions studied.

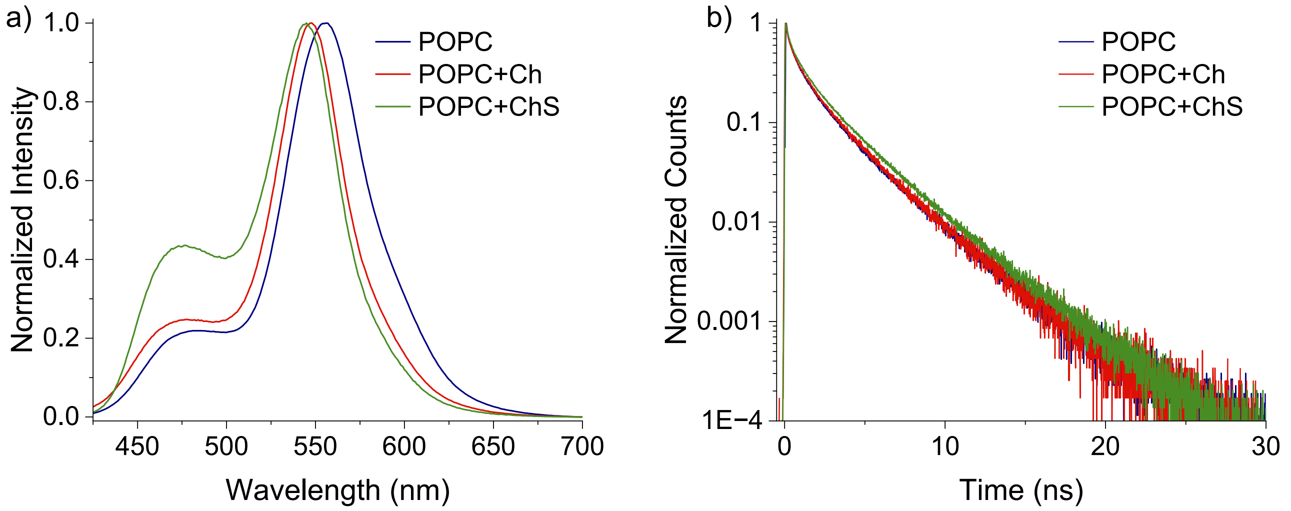


**Figure S4.** (a) Steady-state and (b) Time-resolved fluorescence of $C_{12}$-HPTS (at ROH* position) inside DPPC, DPPC+Ch and DPPC+ChS. The concentrations of Ch and ChS are 20%.

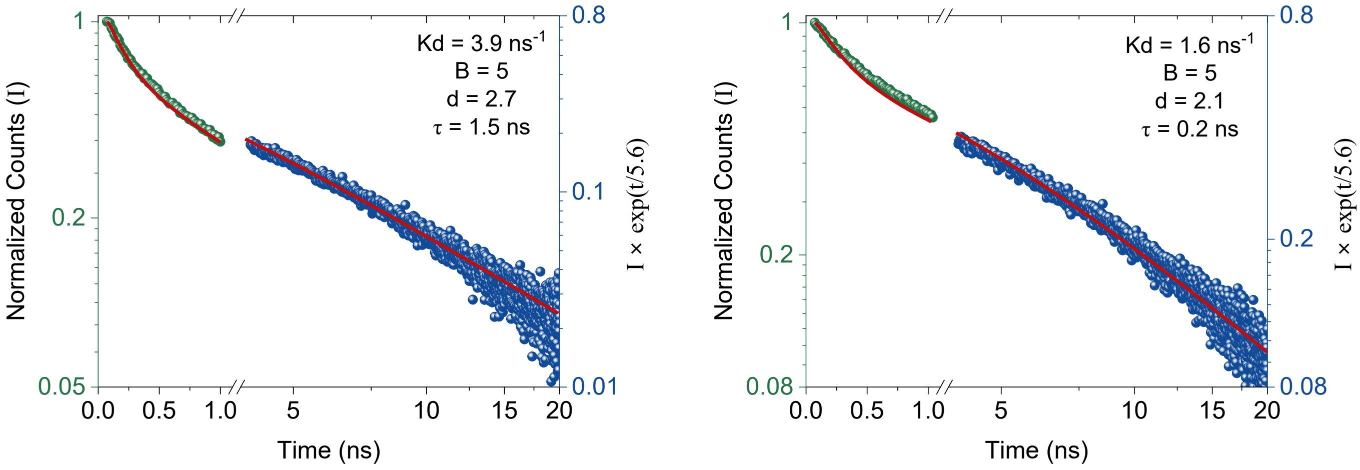


**Figure S5.** Two examples of fitting TCSPC curves: DPPC + 20% Ch (left) and DPPC + 20% ChS (right), where $K_d$ ($k_{PT}$) is extracted from the initial decay, while the dimensionality (d) is extracted from the long-lived tail using Eq.(3).

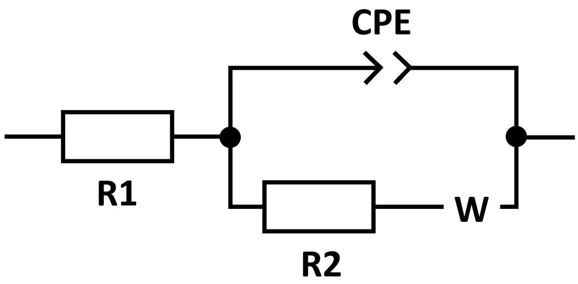


**Figure S6.** Equivalent circuit used for fitting the Nyquist plots. R1 and R2 represent the contact and bulk resistance, respectively. The constant phase element (CPE) describes the capacitance developed as an electrical double layer next to the electrodes and the Warburg element is a diffusion element.
